# Inoculation of indigenous nitrogen-fixers isolated from the Kuwait desert enhances seedling growth and nutrient uptake in a greenhouse bioassay

**DOI:** 10.64898/2025.12.02.691963

**Authors:** Majda K. Suleiman, Ali M. Quoreshi, Anitha J. Manuvel, Mini T. Sivadasan, Sheena Jacob

## Abstract

Desert soil degradation, primarily caused by anthropogenic disturbance and desertification, poses significant challenges for ecosystem restoration in degraded ecosystems. Soil microbial communities, particularly diazotrophs, play a crucial role in different soil system processes, including nutrient cycling, and can improve nitrogen-limited characteristics of nutrient-poor soil systems. Free-living and root-associated nitrogen-fixing bacteria have great potential to enhance nitrogen availability in the nitrogen-limited soil and support host plant growth and nutrient acquisition. Free-living nitrogen-fixing bacteria and root-associated rhizobacteria contribute a substantial amount of nitrogen to ecosystems, including arid lands. This study evaluated the growth performance and nutrient uptake ability of four native plant species of Kuwait inoculated with a consortium of selected indigenous putative diazotrophs isolated from the Kuwait desert. The seedlings of *Vachellia pachyceras* were inoculated with both indigenous root-nodule bacteria isolated from Kuwait desert and a commercial inoculum to evaluate their symbiotic efficiency. The seedlings were cultivated under greenhouse conditions in both native desert soils and in potting mix to assess the extent to which growth medium influenced inoculation response. The primary objective was to determine whether the inoculated indigenous N_2_ fixing bacteria could contribute early seedling development and nutrient acquisition, thereby supporting their potential use them as biofertilizer in future large-scale restoration efforts. Bacterial inoculation significantly enhanced plant dry mass and nutrient uptake across all tested plant species compared to the non-inoculated controls. The magnitude of improvement varied with bacterial density, plant species, and growth medium used.

These findings are consistent with evidence that isolated indigenous N_2_-fixers have the potential to enhance plant growth and nutrient uptake in selected native plant species, supporting their use as biofertilizers for restoration and revegetation efforts in arid environments. This study represents the first evaluation of Kuwait’s native seedlings inoculated with indigenous diazotrophs, highlighting their potential for sustainable ecosystem restoration.

## Introduction

Recent land degradation in the Kuwait desert, resulting from prevailing environmental extremes and anthropogenic disturbances, is a significant environmental concern that contributes to further desertification and poor soil conditions. Consequently, degradation of native vegetation communities affects the characteristics of desert vegetation. Nevertheless, desert plants are indigenous plants adapted to the extreme local environment. They are characterized by slower growth and adaptation mechanisms that enable them to survive in harsh environments. However, it is extremely challenging to succeed in natural regeneration and revegetation in degraded soil conditions, as the soils have already lost their essential nutrient content, organic matter, and key biological soil properties, including the soil microbial community [1]. Evidences suggest that the native plants’ population in Kuwait has been diminishing in recent years due to soil degradation, extreme weather, climatic fluctuations and various anthropogenic activities [2, 3]. Restoration and revegetation of the native plants are essential to protect them from extinction. To restore the native plant community successfully, it is important to recover soil fertility and the rhizosphere microbial community, as the soil is the major source of nutrients for plant growth and development. Nevertheless, desert lands host diverse resilient microbial communities that exist as free-living, associate with plants, or form soil biocrusts [4].

The soil microbial communities are considered as the key driving force for regulating critical soil ecosystems processes, such as nutrient cycling and decomposition of organic matter, nutrient mineralization, and biogeochemical cycling of terrestrial ecosystems [5, 6, 7, 8]. The available soil nutrients for native plant are deficient in Kuwait’s desert soil due to lack in organic matter (<1%), low clay materials, high in calcareous materials, and low in essential nutrients [9]. Availability of soil nutrients is one of the important factors for the restoration of the degraded desert soil. The extreme arid environmental conditions generally recognized that nutrient biogeochemical cycling, especially N cycling are microbially mediated [10]. Soil microbial community including N fixing bacteria could play an important role in nutrient cycling, decomposition of organic matter, and improvement of soil fertility, and maintenance of healthy ecosystems [11]. To plan effective strategies for conserving and restoring desert ecosystems, it is often necessary to manipulate and establish associated soil microorganisms for the successful establishment of vegetation cover.

Diazotrophic bacteria are a group of bacteria that dwell in the rhizospheric soil or are associated with the root system of plants and fix atmospheric dinitrogen to plant-available nutrients. These diazotrophs have great potential to alleviate nitrogen scarcity in nitrogen-scarce soils and enhance the accessibility of nitrogen to the host plant [12, 13, 14].

Bashan et al. [15] and Moreno et al. [16] demonstrated that the PGPB enhanced the growth of native plant species and improved soil stability in degraded forest and desert soils. Liu et al. [17] reported that these bacteria accelerated plant development and improved the survival rate of different forest seedlings. Inoculation of plant-growth-promoting (PGP) nitrogen-fixing bacteria is an alternative to the use of chemical fertilizers [18]. The generous use of chemical fertilizers triggers environmental concern and economic losses. In view of the growing interest in developing alternative approaches to improve soil fertility, the use of bioinoculants and biofertilizers containing plant-beneficial microorganisms represents a promising alternative to chemical fertilizers. Bioinoculants have the potential to mobilize the essential nutrients to the plants [19]. Deficiency of available nitrogen to plants in the Kuwait’s desert soil signifies the necessity to develop a strategy to enhance its availability via diazotrophic bacteria to improve the soil fertility [20]. Although numerous studies have documented the significant impact of plant growth-promoting bacteria (PGPB) and other beneficial microbes on crop yield and productivity, limited research has focused on their potential to enhance the growth and biomass of desert seedlings/plants through free-living or root-associated beneficial microorganisms [21].

Application of indigenous free-living diazotrophs and root-associated plant growth – promoting bacteria for promoting the growth of Kuwait native plants remains unexplored. In this current study, we evaluated the plant growth performance and nutrient uptake after inoculation of free-living nitrogen-fixing bacterial isolates and root-associated bacteria, obtained from our previous study [20] to *Rhanterium epapposum, Farsetia aegyptia, Haloxylon, Vachellia pachyceras Haloxylon salicornicum* under greenhouse condition. The hypothesis of this research was that inoculation of free-living and root-associated diazotrophs may increase growth and nutrient uptake of native desert plants under the current experimental setting. Therefore, our main objective of this study was to evaluate the potential of indigenous nitrogen-fixing bacteria isolates from Kuwait’s desert soils to enhance seedling growth performance and nutrient uptake of native desert plants under controlled greenhouse conditions.

## Materials and methods

### Inoculum preparation

The bacterial isolates used in this greenhouse experiment are indigenous diazotrophic strains isolated from the rhizospheric soil of *Rhanterium epapposum*, *Farsetia aegyptia*, and *Haloxylon salicornicum* (free-living nitrogen-fixing bacteria), as well as root nodules of *Vachellia pachyceras* (symbiotic nitrogen-fixers). The nitrogen-fixing capacity of isolated diazotrophs was previously tested and confirmed using the Acetylene Reduction Assay, and they were identified using the 16s rRNA gene sequencing method [20]. Totally, 6, 3, 11 and 19 strain of indigenous nitrogen-fixers were used as inoculum to test on the native plant species *Rhanterium epapposum, Farsetia aegyptia, Haloxylon salicornicum* and *Vachellia pachyceras*, respectively. For each bacterial isolate obtained from a single plant species, cell suspensions were prepared at a density of 10^8^ CFU mL^-1^ [22, 23]. These isolates were then pooled together to form a mixture of bacterial inoculum with a final cell density of 10^8^ CFU mL^-1^. *From this mixture, a single dilution of 10⁴ CFU mL⁻¹ was prepared for Farsetia aegyptia, Rhanterium epapposum*, and *Haloxylon salicornicum* (free-living nitrogen-fixing bacteria). For Vachellia pachyceras, a dilution series of 10⁸, 10⁶, 10⁴, and 10² CFU mL⁻¹ was prepared.

### Seedling Production

Two types of seedling growth media (desert soil and potting mix) were used in the experiment. Desert soil was collected from KSRI, packed in double autoclavable bags, and transported to the laboratory. Potting mix soil was prepared by mixing agricultural soil, peat moss, and potting soil in a 2:1:1 volume-to-volume ratio (v/v) and packed into double autoclavable bags. The packed soils were sterilized twice in an autoclave at 121°C for 30 minutes, with complete cooling and mixing between cycles, and then air-dried. Likewise, jiffy pots and potting mix soil used in the initial seedling production were also sterilized prior to use. The irrigation water was sterilized once in an autoclave at 121°C for 15 min and used throughout the experiment. The sterilized jiffy pots were filled with autoclaved potting mix soil or desert soil and saturated with sterile water, and placed in a trey (30 jiffy pots). As Kuwait’s native plant seeds were very sensitive to chemical treatment, the surface sterilization was done by soaking *Rhanterium epapposum* and *Farsetia aegyptia* overnight in sterile water and washed six times in sterile water, and one capitulum of *Rhanterium epapposum* was sown on the surface of the sterile potting mix soil or desert soil in jiffy pots. The seeds of *Farsetia aegyptia* were sown in jiffy pots filled with the potting mix or desert soil for germination. *Haloxylon salicornicum* seeds were washed six times with sterile water, and one seed was sowed on the top of the potting mix soil in jiffy pot for germination. *Vachellia pachyceras* seeds were treated with concentrated sulphuric acid for 30 min [24], washed six times in sterile water, and soaked in sterile water for overnight. The imbibed seeds were transferred to sterile petri-plate with moistened filter paper until germination. The germinated seeds were transferred to sterile jiffy pots with potting soil mix or desert sand. All the trays were maintained in the standard greenhouse conditions and irrigated with sterile water throughout the experiment as required. Transplantation and Inoculation Two-weeks after the germination in the jiffy pot, the seedlings of all the species were transplanted to one-gallon pots and maintained in the greenhouse. Six-weeks after the transplantation, the seedlings were inoculated with 20 ml of indigenous bacterial suspension with desired cell densities (10^4^ and 10^8^ CFU mL^-1^ for *Rhanterium epapposum, Farsetia aegyptia, Haloxylon salicornicum* whereas 10^2^, 10^4^, 10^6^ 10^8^ CFU mL^-1^ for *Vachellia pachyceras*) [25, 26]. Seedlings of all the species in the control treatment were not inoculated. Each treatment including control had eight seedlings as replicates which were arranged in Completely Randomized Design (CRD) in the greenhouse conditions. Furthermore, *Vachellia pachyceras* inoculated with 10 ml of 1 OD commercial American Type Culture Collection (ATCC) *Rhizobium leguminosarum* bv. viceae (10004; LOT: 59679216) and *Bradyrhizobium* sp. (BAA-1182; LOT: 60865886) in triplicate were also prepared and maintained along with the main experiment, to compare the potential of commercial inoculum to form nodulation and improve nitrogen uptake of *Vachellia pachyceras* under greenhouse conditions.

### Data Collection and Data Analysis

The inoculated and non-inoculated control seedlings were maintained under greenhouse conditions, and their growth performance (plant height, number of leaves, stem diameter) and physical condition (vigor) were recorded before the experiment was terminated after the 10^th^ month. At the end of the experiment, five randomly selected seedlings per treatment were excavated completely, and the soil was rinsed off the roots. The number and weight of the root nodules of *Vachellia pachyceras* were recorded. The seedlings were separated into roots and shoots and dried in an oven at 70°C for 48 hr or until a constant dry weight was achieved. The dry mass of roots and shoots of each plant species was determined on an analytical scale. The dried powdered shoot samples of each plant species were analyzed for Nitrogen (N), Phosphorus (P), and Potassium (K).

Seedling biomass, plant parameters and chemical analysis data were analyzed using Analysis of Variance Procedure (ANOVA) using SPSS® software - version 22 (IBM®) and means were compared using the Duncan’s Multiple Range test to ascertain the significant differences among treatments at the 5% level of probability [27]. Type soil medium and bacterial inoculum were considered as factors in the analysis.

## Results

Effect of potential nitrogen-fixer bacterial inoculations on plant growth parameters of selected native plant species

### Rhanterium epapposum

Generally, inoculation with the two densities (10^4^ and 10^8^) of indigenous bacterial inoculum to *Rhanterium epapposum* significantly enhanced the average root (p <0.001), shoot (p <0.001), and total biomass (p <0.001) per plant compared to the non-inoculated control (Table 1). Additionally, average root biomass per plant increased significantly (p <0.001) in plants grown in potting soil when compared to that grown in desert soil (Table 1). However, bacterial inoculation and effect of growth medium types used had no significant impact on any other growth parameters determined when compared to the control (Table 1). The average root biomass of the seedlings grown in desert soil medium increased by 88% when compared to the seedlings grown in potting mix soil (Fig 1). Likewise, the indigenous bacterial inoculation increased average root biomass by 331% and 538%, average shoot biomass by 198% and 303% and average total biomass by 206% and 318% when inoculated with 10^4^ CFU/ml and 10^8^ CFU/ml densities (Fig 1). Although there was significant difference in the average root biomass and shoot biomass between non-inoculated control and those inoculated with indigenous bacteria, there was no significant difference within the two inoculum densities used (10⁴ CFU/ml and 10⁸ CFU/ml) (Fig 1). Unlike the average root and shoot biomass, the average total biomass was significantly affected by inoculum density, which resulted in 54% increase in average total biomass of seedlings inoculated with indigenous inoculum of density 10^8^ cells compared to those inoculated with cell density of 10^4^ (Fig 1).

**Fig 1.**
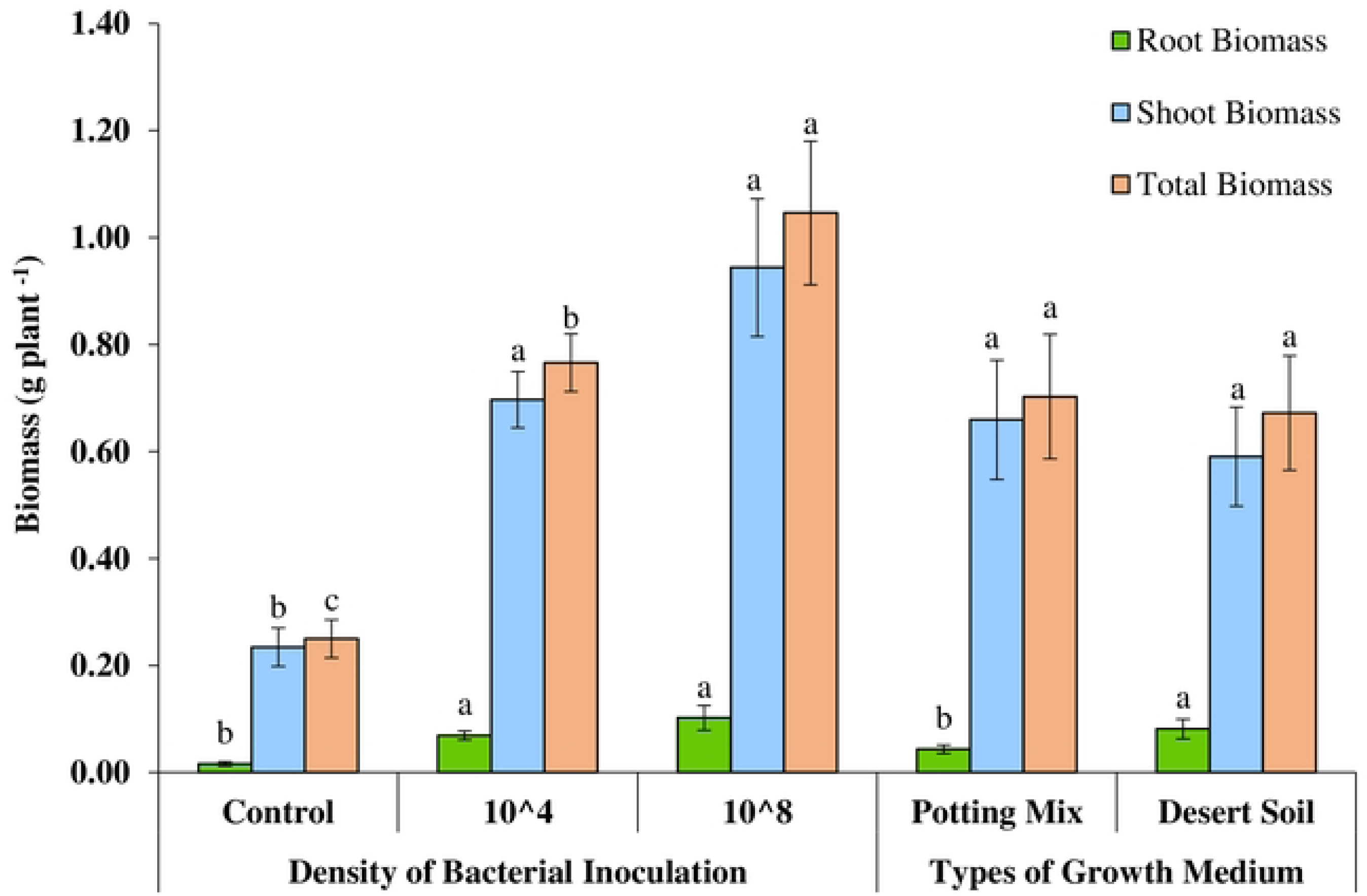
Effect of type of growth medium and density of bacterial inoculation on biomass per plant in *Rhanterium epapposum*.

**Table 1.** Results of Analysis of Variance (P values) Testing for the Effect of Growth medium and Bacterial Inoculum on Growth Parameters of Selected Native Plant Species.

| Variable | Plant Height | Stem Diameter | Plant Vigor | Root Biomass | Shoot Biomass | Total Biomass | Number of Root Nodules | Weight of Nodules | Root Shoot Ratio |
| --- | --- | --- | --- | --- | --- | --- | --- | --- | --- |
| *Rhanterium epapposum* |  |  |  |  |  |  |  |  |  |
| Soil Media | 0.499 | 0.386 | 0.584 | 0.011 | 0.496 | 0.769 | NA | NA | 0.150 |
| Bacterial Inoculation | 0.118 | 0.285 | 0.510 | <0.001 | <0.001 | <0.001 | NA | NA | 0.894 |
| Soil Media * Bacterial Inoculation | 0.243 | 0.413 | 0.114 | 0.054 | 0.725 | 0.677 | NA | NA | 0.585 |
| *Farsetia aegyptia* |  |  |  |  |  |  |  |  |  |
| Soil Media | 0.016 | 0.026 | 0.170 | 0.065 | 0.001 | <0.001 | NA | NA | 0.546 |
| Bacterial Inoculation | 0.453 | 0.795 | 0.613 | 0.005 | <0.001 | <0.001 | NA | NA | 0.244 |
| Soil Media * Bacterial Inoculation | 0.194 | 0.413 | 0.613 | 0.987 | 0.335 | 0.332 | NA | NA | 0.872 |
| *Haloxylon salicornicum* |  |  |  |  |  |  |  |  |  |
| Soil Media | 0.082 | 0.646 | 1.000 | 0.007 | 0.001 | <0.001 | NA | NA | 0.009 |
| Bacterial Inoculation | 0.373 | 0.653 | 0.613 | 0.416 | <0.001 | <0.001 | NA | NA | 0.961 |
| Soil Media * Bacterial Inoculation | 0.138 | 0.414 | 0.243 | 0.428 | 0.046 | 0.050 | NA | NA | 0.907 |
| *Vachellia pachyceras* |  |  |  |  |  |  |  |  |  |
| Soil Media | 0.653 | 0.366 | 0.057 | 0.893 | 0.481 | 0.787 | 0.806 | 0.934 | 0.200 |
| Bacterial Inoculation | 0.114 | 0.058 | 0.139 | 0.001 | 0.002 | <0.001 | 0.009 | 0.010 | 0.131 |
| Soil Media * Bacterial Inoculation | 0.090 | 0.084 | 0.139 | 0.195 | 0.037 | 0.031 | 0.084 | 0.007 | 0.784 |

### Farsetia aegyptia

*Farsetia aegyptia*, seedlings inoculated with the indigenous bacterial inoculum at 10⁸ cells exhibited significantly greater root (P ≤ 0.005), shoot (P ≤ 0.001), and total biomass (P ≤ 0.001) compared to those inoculated with 10⁴ cells and the control (Table 1). Bacterial inoculum did not have any significant effect on other growth parameters.

Interestingly, plant height (P ≤ 0.016), stem diameter (P ≤ 0.026), shoot biomass (P≤0.001) and total biomass (P<0.001) of seedlings grown in desert soil was significantly higher than those grown in potting mix. Soil medium did not have significant effect on other growth parameters (Table 1).

Nevertheless, the average plant height, stem diameter, shoot biomass, and total biomass of *Farsetia aegyptia* seedlings grown in desert soil increased by 26%, 24%, 58%, and 58%, respectively, compared to seedlings grown in the potting-mix medium (Fig 2). A significant increase in shoot biomass (177%) and total biomass (182%) was observed in seedlings inoculated with the indigenous bacterial inoculum at a density of 10⁸ CFU/ml cells compared to the control (Fig 2). Although inoculation with the 10⁴ CFU/ml cell density also resulted in increases in these parameters, the improvements were not statistically significant relative to the control plants.

**Fig 2.**
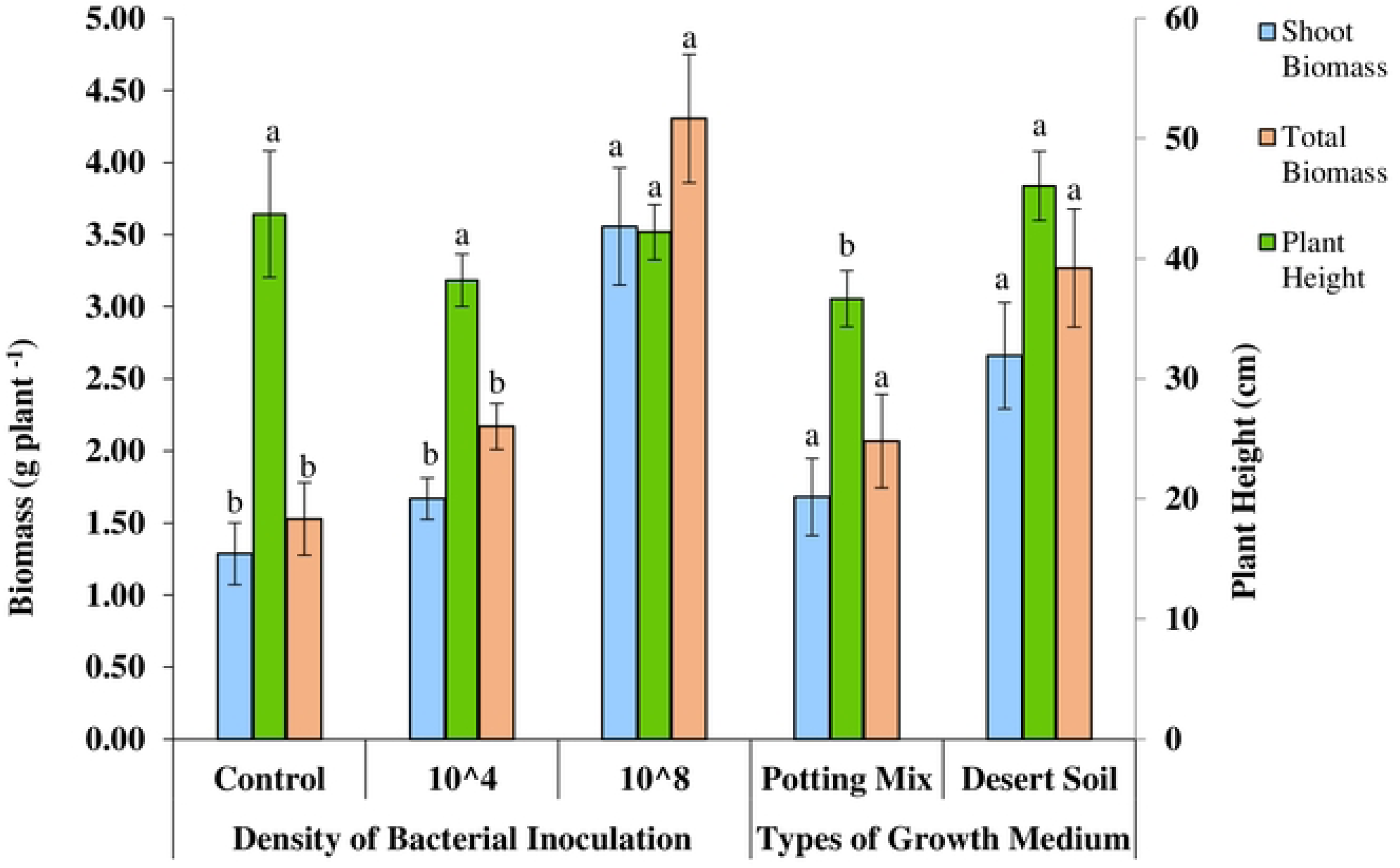
Effect of type of growth medium and density of bacterial inoculation on biomass per plant, plant height and stem diameter of *Farsetia aegyptia*.

### Haloxylon salicornicum

Interestingly, the seedling growth substrate had a remarkable effect, significantly increased the average root biomass (P ≤ 0.007), shoot biomass (P ≤ 0.001), total biomass (P < 0.001), and root-to-shoot ratio (P ≤ 0.009) of *Haloxylon salicornicum* seedlings (Table 1). Bacterial inoculation at both 10⁴ CFU/ml and 10⁸ CFU/ml cell densities also significantly increased shoot biomass (P ≤ 0.001), and total biomass (P ≤ 0.001), compared to the non-inoculated control (Table 1). However, neither growth substrate nor bacterial inoculation significantly affected the other measured growth parameters compared to the control (Table 1).

The average root biomass, shoot biomass, and total biomass of *Haloxylon salicornicum* seedlings grown in desert soil increased by 476%, 46%, and 75%, respectively, compared with seedlings grown in the potting-mix medium (Fig 3). The average shoot biomass significantly increased by 89% and 123% in seedlings inoculated with indigenous bacterial inoculum at densities of 10⁴ CFU/ml and 10⁸ CFU/ml cells, respectively (Fig 3). Similarly, the average total biomass significantly increased by 95% and 123% in seedlings inoculated with 10⁴ CFU/ml and 10⁸ CFU/ml cell densities, respectively (Fig 3). However, there was no significant difference between the two inoculum densities with respect to shoot or total biomass.

**Fig 3.**
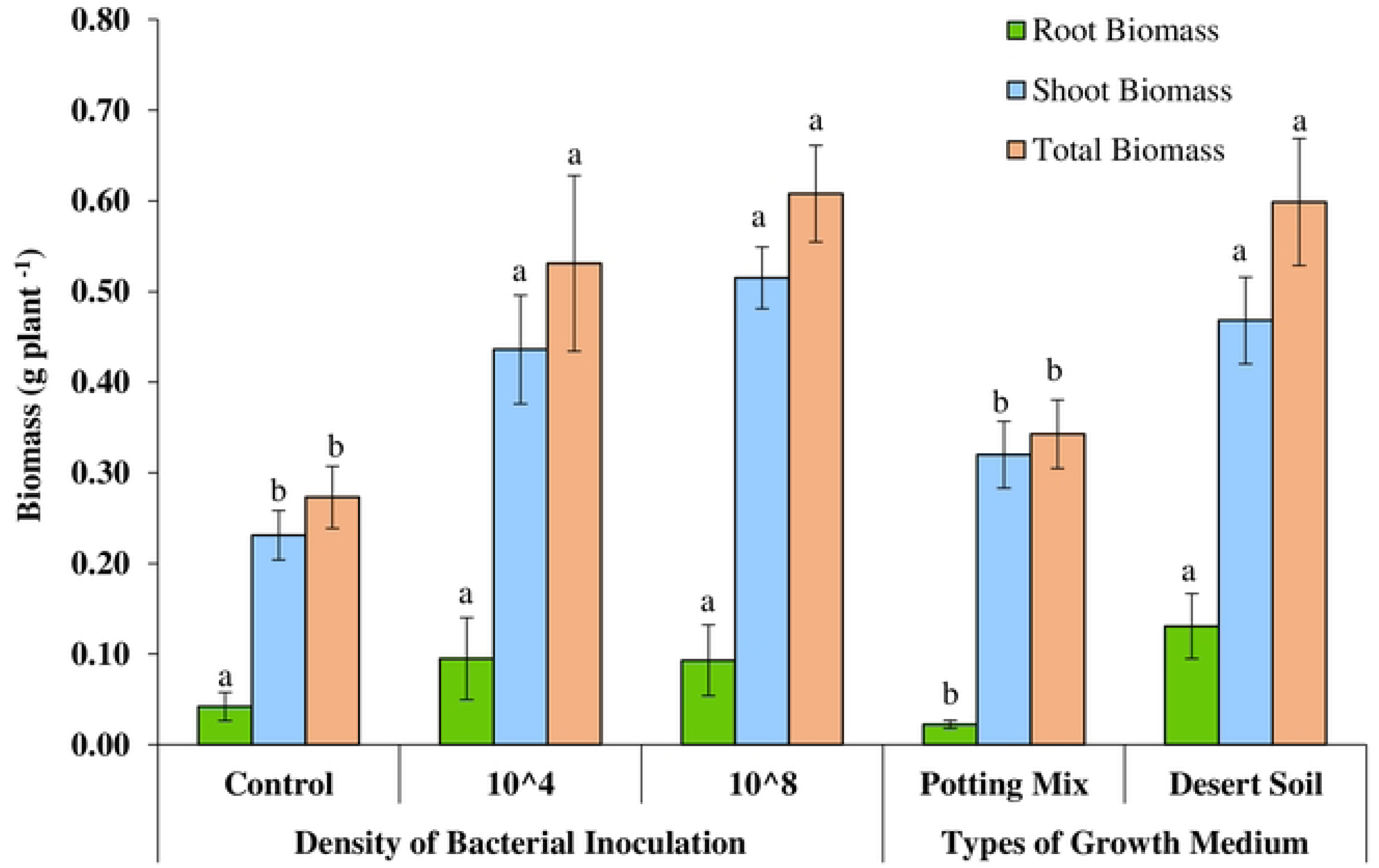
Effect of type of growth medium and density of bacterial inoculation on biomass per plant of *Haloxylon salicornicum*.

### Vachellia pachyceras

There was a significant increase in average root biomass (P ≤ 0.001), shoot biomass (P ≤ 0.002), total biomass (P ≤ 0.001), number of root nodules (P ≤ 0.009), and dry weight of root nodules (P ≤ 0.010) in seedlings inoculated with the indigenous bacterial inoculum, with the 10² CFU/ml and 10⁴ CFU/ml cell densities producing the most effective responses out of the four densities used i.e. 10^2^, 10^4^, 10^6^ and 10^8^ CFU/ml cells (Table 1 and Fig 4). However, neither inoculation nor growth media had a significant influence on the other recorded growth parameters (Fig 4).

**Fig 4.**
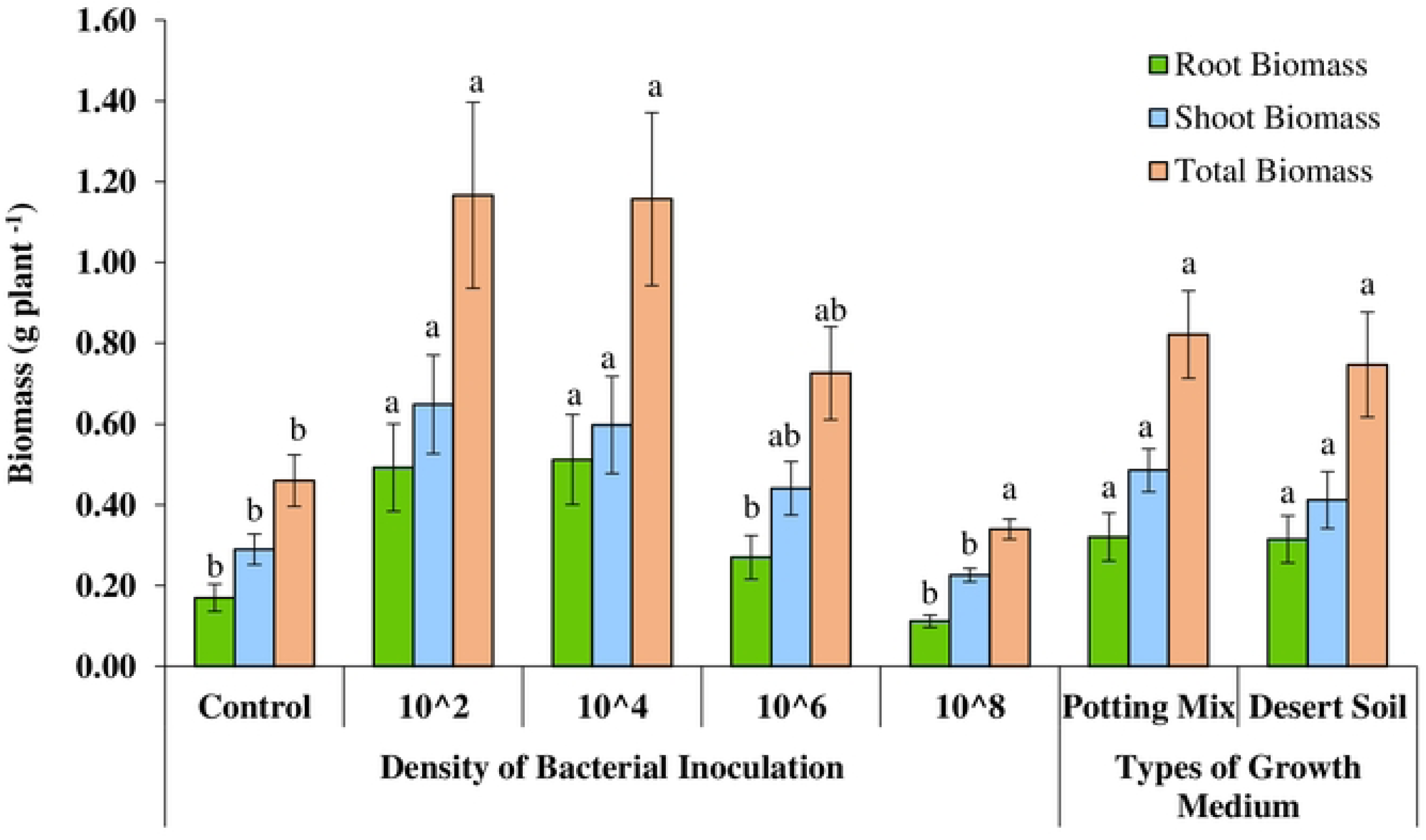
Effect of type of growth medium and density of bacterial inoculation on biomass per plant of *Vachellia pachyceras*.

The average root biomass increased by 190%, and 201% in seedlings inoculated with indigenous bacterial inoculum at densities of 10² CFU/ml and 10⁴ CFU/ml cells, respectively (Fig 4). Similarly, the average shoot biomass increased by 124% and 106% the total biomass increased by 154% and 152% in seedlings inoculated with 10² CFU/ml and 10⁴ CFU/ml cell densities, respectively (Fig 4). Interestingly, the highest concentration of indigenous bacterial inoculum (at 10^8^ CFU/ml density) had no significant increase on average root biomass, shoot biomass and total biomass of the seedlings when compared to the non-inoculated control (Fig 4). The average number and dry weight of root nodules had significant effect in seedlings inoculated with indigenous bacterial inoculum at densities 10^2^, 10^4^ cells.

Effect of putative nitrogen-fixer bacterial inoculation on plant nutrient uptake

In general, inoculation of putative nitrogen-fixing bacteria at various densities significantly affected the nutrient uptake of all four selected native plant species compared to the non-inoculated control (Table 2) under greenhouse experimental conditions. However, the increase in nutrient uptake ability of the tested bacterial inoculum varied depending on the density of bacteria in the inoculum used. In contrast, there was no significant interaction effect of growing media and inoculation on the nutrient uptake except in *Farsetia aegyptia* and *Haloxylon salicornicum* (Table 2).

**Table 2.** Results of Analysis of Variance (P values) Testing for the Effect seedling growth Medium and Bacterial Inoculation on N, P, K, Mg, Na, Ca, C and S uptake on Selected Native Plant Species.

| Variable | N | P | K | Mg | Na | Ca | C | S |
| --- | --- | --- | --- | --- | --- | --- | --- | --- |
| *Rhanterium epapposum* |  |  |  |  |  |  |  |  |
| Soil Type | 0.573 | 0.102 | 0.342 | 0.531 | 0.160 | 0.056 | 0.717 | 0.988 |
| Bacterial Inoculation | <0.001 | 0.001 | <0.001 | <0.001 | 0.001 | 0.001 | <0.001 | <0.001 |
| Soil Type * Bacterial Inoculation | 0.532 | 0.308 | 0.700 | 0.905 | 0.538 | 0.554 | 0.367 | 0.709 |
| *Farsetia aegyptia* |  |  |  |  |  |  |  |  |
| Soil Type | <0.001 | 0.189 | 0.010 | 0.463 | 0.655 | 0.011 | 0.013 | 0.004 |
| Bacterial Inoculation | <0.001 | <0.001 | 0.001 | 0.002 | 0.105 | 0.001 | 0.001 | 0.001 |
| Soil Type * Bacterial Inoculation | 0.181 | 0.197 | 0.126 | 0.221 | 0.062 | 0.372 | 0.540 | 0.167 |
| *Haloxylon salicornicum* |  |  |  |  |  |  |  |  |
| Soil Type | 0.002 | 0.001 | 0.494 | 0.084 | 0.051 | 0.027 | 0.007 | 0.272 |
| Bacterial Inoculation | 0.014 | 0.023 | 0.002 | 0.001 | 0.007 | 0.001 | 0.004 | 0.011 |
| Soil Type * Bacterial Inoculation | 0.321 | 0.180 | 0.264 | 0.048 | 0.541 | 0.073 | 0.138 | 0.089 |
| *Vachellia pachyceras* |  |  |  |  |  |  |  |  |
| Soil Type | 0.647 | 0.804 | 0.397 | 0.275 | 0.051 | 0.308 | 0.910 | 0.313 |
| Bacterial Inoculation | 0.022 | 0.042 | 0.032 | 0.015 | 0.039 | 0.029 | 0.026 | 0.040 |
| Soil Type * Bacterial Inoculation | 0.125 | 0.186 | 0.292 | 0.506 | 0.086 | 0.229 | 0.198 | 0.131 |

N: Nitrogen; P: Phosphorus; K: Potassium; Mg: Magnesium; Na: Sodium; Ca: Calcium; C:
Carbon; S: Sulphur.

### Rhanterium epapposum

Inoculation of *Rhanterium epapposum* with two densities (10⁴ CFU/ml and 10⁸ CFU/ml) of putative nitrogen-fixing bacterial inoculum in the rhizosphere significantly increased shoot nitrogen (N) (P ≤ 0.001), phosphorus (P) (P ≤ 0.001), and potassium (K) (P ≤ 0.001) content compared to the non-inoculated control (Table 2). Specifically, shoot N content increased by 176% and 319%, shoot P content by 185% and 244%, and shoot K content by 215% and 358% when inoculated with 10⁴ CFU/ml and 10⁸ CFU/ml cell densities, respectively, using the mixture of isolated indigenous putative nitrogen-fixers (Fig 5). The cell density of 10^8^ CFU/ml produced the best results.

**Fig 5.**
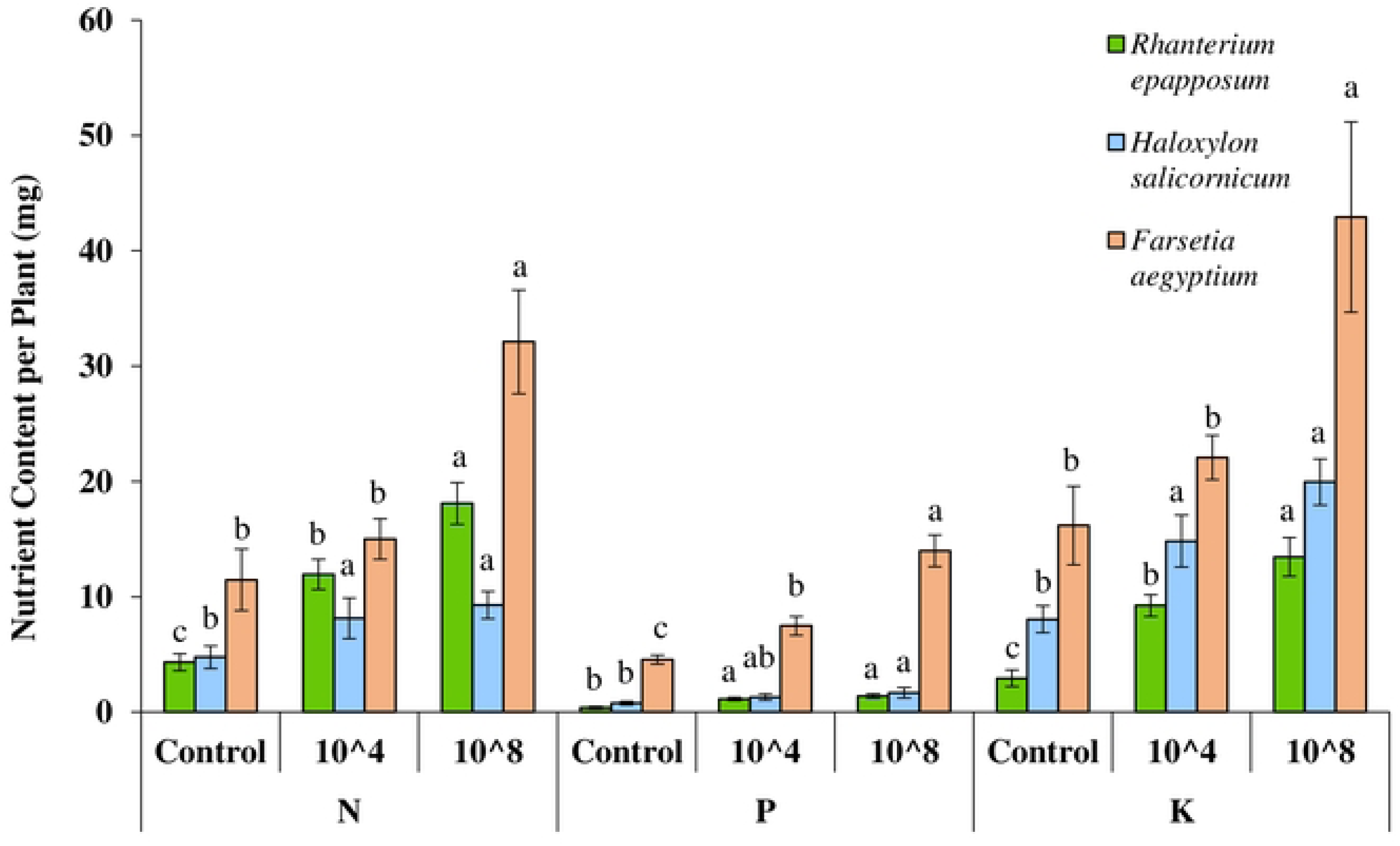
Effect of bacterial inoculation on N, P, K uptake measured as shoot N, P, K content per plant in *Rhanterium epapposum, Haloxylon salicornicum and Farsetia aegyptia*.

Additionally, inoculation also significantly enhanced shoot magnesium (Mg) (P ≤ 0.001), sodium (Na) (P ≤ 0.001), calcium (Ca) (P ≤ 0.001), carbon (C) (P ≤ 0.001), and sulfur (S) (P ≤ 0.001) content (Table 2). However, neither the type of growth medium nor its interaction with bacterial inoculation had a significant effect on nutrient uptake (Table 2).

### Farsetia aegyptia

Inoculation of *Farsetia aegyptia* with putative nitrogen-fixing bacterial inoculum in the rhizosphere significantly increased shoot nitrogen (N) (P ≤ 0.001), phosphorus (P) (P ≤ 0.001), and potassium (K) (P ≤ 0.001) uptake compared to the non-inoculated control (Table 2). Inoculation with the 10⁸ cell density resulted in significant increases in shoot N, P, and K by 180%, 207%, and 165%, respectively (Fig 5). Furthermore, inoculation also significantly enhanced shoot uptake of magnesium (Mg) (P ≤ 0.002), calcium (Ca) (P ≤ 0.001), carbon (C) (P ≤ 0.001), and sulfur (S) (P ≤ 0.001) (Table 2).

Interestingly, the seedling growth media also had a significant effect on shoot nutrient status, influencing N (P ≤ 0.001), K (P ≤ 0.010), Ca (P ≤ 0.011), C (P ≤ 0.013), and S (P ≤ 0.004) content of the plant shoot (Table 2). However, the growth media had no significant effect on other parameters tested, including shoot P, Mg, and Na content (Table 2). *Haloxylon salicornicum* A similar trend was observed in *Haloxylon salicornicum* as the above mentioned native plant species. Shoot nitrogen (N) (P ≤ 0.014), phosphorus (P) (P ≤ 0.023), and potassium (K) (P ≤ 0.002) content increased significantly in seedlings inoculated with the indigenous bacterial inoculum compared to the non-inoculated control (Table 2). As shown in Figure 5, inoculation increased shoot N content by 70% and 95%, shoot P content by 68% and 115%, and shoot K content by 84% and 148% when inoculated with 10⁴ CFU/ml and 10⁸ CFU/ml cell densities, respectively. There was no significant difference between the effects of 10⁴ and 10⁸ cell densities on the shoot N and K content. In addition to N, P, and K, bacterial inoculation also significantly enhanced shoot Mg (P ≤ 0.001), Na (P ≤ 0.007), Ca (P ≤ 0.001), C (P ≤ 0.004), and S (P ≤ 0.011) content (Table 2).

The growth media had a significant positive effect on several nutrient parameters. Seedlings grown in desert soil showed higher shoot N (P ≤ 0.002), Ca (P ≤ 0.027), and C (P ≤ 0.007) contents (Table 2), whereas those grown in potting soil mix recorded significantly higher shoot P (P < 0.001). In contrast, the growth media had no significant influence on shoot K, Mg, Na, or S contents (Table 2).

### Vachellia pachyceras

Inoculation of *Vachellia pachyceras* seedlings with different densities of indigenous bacterial inoculum resulted in significant increases in shoot N (P ≤ 0.022), P (P ≤ 0.042), K (P ≤ 0.032), Mg (P ≤ 0.015), Na (P ≤ 0.039), Ca (P ≤ 0.029), C (P ≤ 0.026), and S (P ≤ 0.040) contents (Table 2). Among the inoculum densities, the 10² CFU/ml level produced the greatest enhancement, increasing shoot N, P, and K contents by 152%, 129%, and 193%, respectively (Fig 6). Similar trends observed in shoot N, P, and K content and increased by 136%, 161%, and 141%, respectively when inoculated with 10^4^ CFU/ml density of bacterial inoculum (Fig 6). No significant effects of growth media or their interaction with bacterial inoculation were observed for any of the measured shoot nutrients (Table 2).

**Fig 6.**
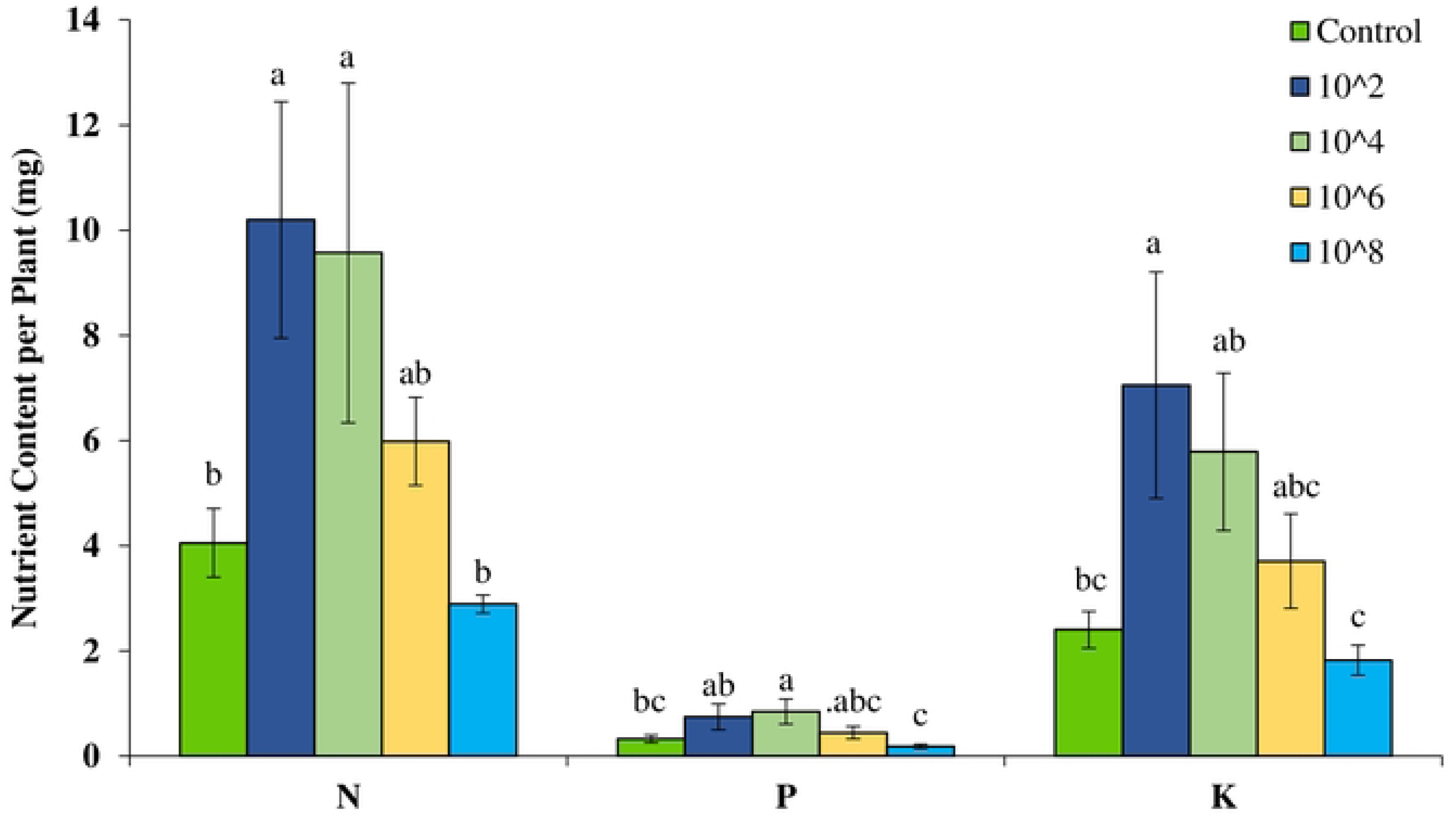
Effect of bacterial inoculation on N, P, K uptake measured as shoot N, P, K content per plant in *Vachellia pachyceras*.

Evaluation of Commercial Inoculum on *Vachellia pachyceras*

The growth and nutrient uptake of *V. pachyceras* were also evaluated, after inoculation with the commercial ATCC bacterial strains, following the same procedure used for the indigenous bacterial inoculum experiments. The results are presented in Table 3.

**Table 3.** Results of Analysis of Variance (P values) Testing for the Effect of Commercial Inoculum on Nodulation, Growth Parameters, and Nitrogen Content of *Vachellia pachyceras*.

| Variable | Number of Nodules | Plant Height | Shoot Biomass | N Content |
| --- | --- | --- | --- | --- |
| Soil Media | 0.009 | 0.001 | 0.002 | 0.001 |
| Inoculation | 0.091 | 0.114 | 0.022 | 0.017 |
| Soil Media * Inoculation | 0.187 | 0.121 | 0.355 | 0.347 |

The seedlings grown in potting soil mix significantly increased average plant height by 75%, (P ≤ 0.001), shoot biomass by 197%, (P ≤ 0.002), number of nodules by 394% (P ≤ 0.009), and nitrogen content by 247% (P ≤ 0.001) compared to those grown in desert soil (Fig 7 and 8). The *Vachellia pachyceras* seedlings, when inoculated with commercial *Rhizobium leguminosarum* ATCC and *Bradyrhizobium* sp. ATCC significantly increased the average shoot biomass (P ≤ 0.022) and nitrogen (P ≤ 0.017) content when compared to the non-inoculated control.

**Fig 7.**
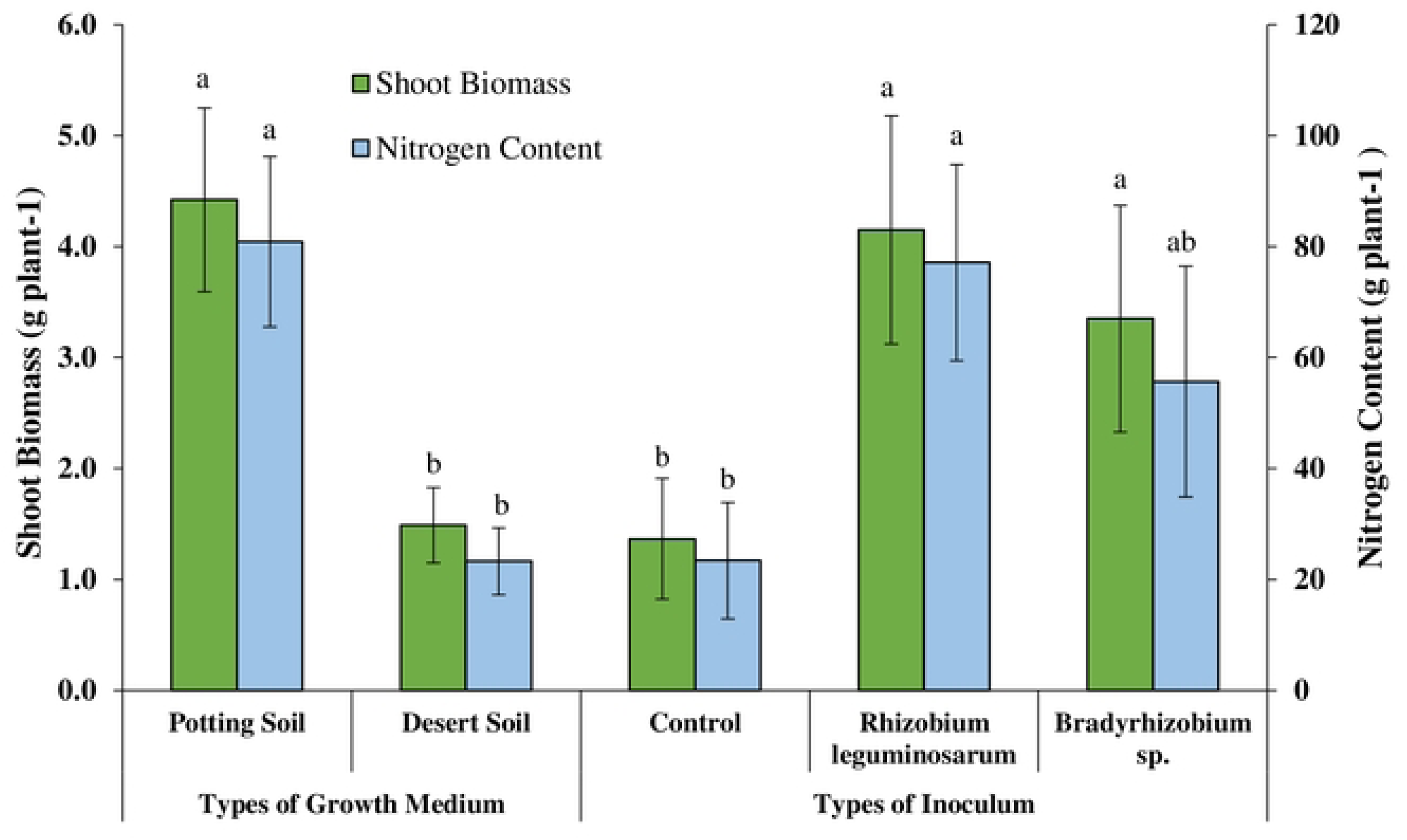
Effect of type of growth medium and commercial inoculum on shoot biomass and nitrogen content per plant of *Vachellia pachyceras*.

**Fig 8.**
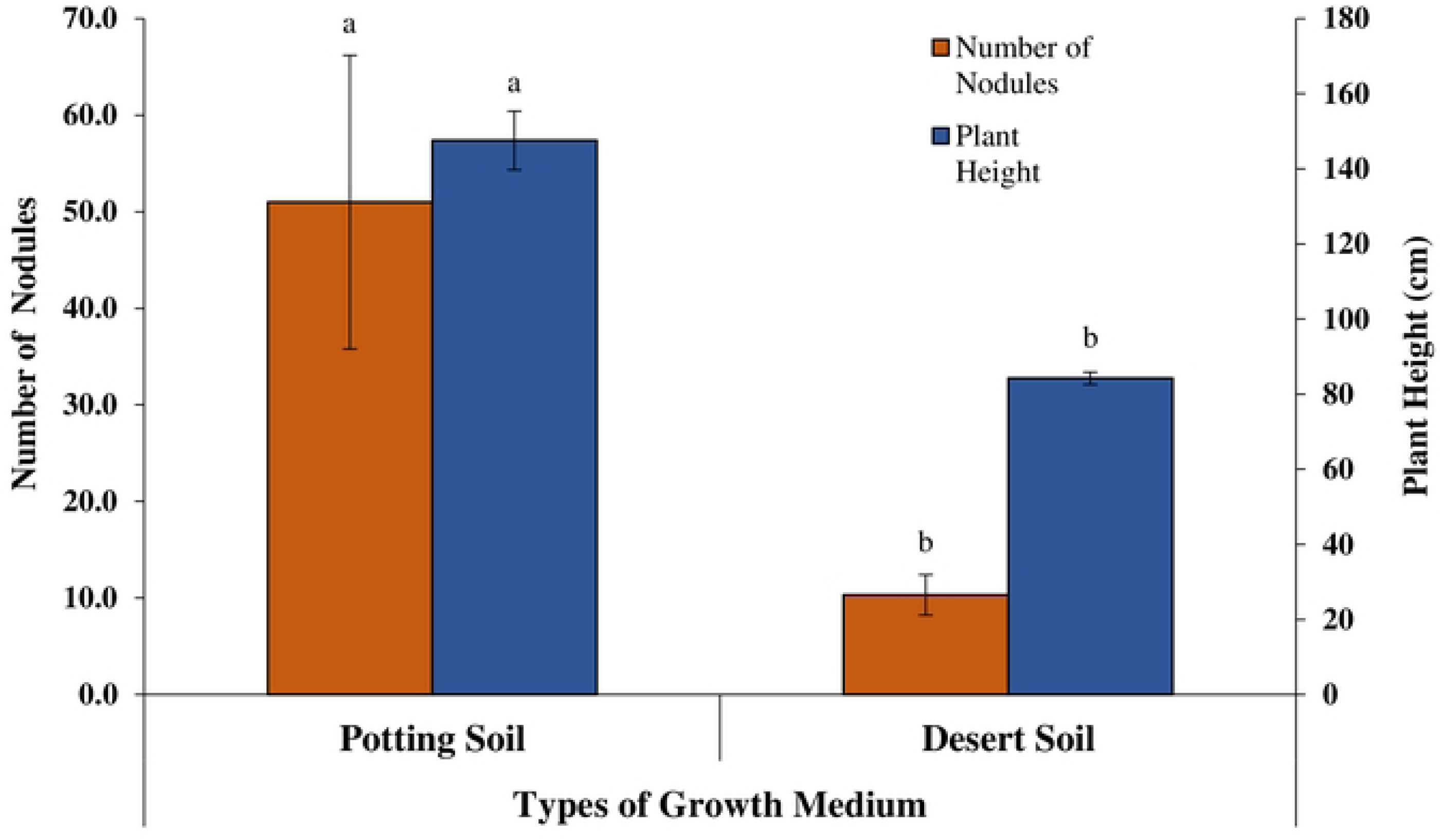
Effect of type of growth medium and commercial inoculum on number of nodules and plant height of *Vachellia pachyceras*.

The average shoot biomass of the *Vachellia pachyceras* seedlings significantly increased by 204% and 145% when inoculated with *Rhizobium leguminosarum* ATCC and *Bradyrhizobium* sp. ATCC, respectively, when compared to the non-inoculated control (Fig 7. Likewise, the average nitrogen content of *Vachellia pachyceras* seedlings significantly increased by 229% and 138% when inoculated with *Rhizobium leguminosarum* ATCC and *Bradyrhizobium* sp. ATCC, respectively, compared to the non-inoculated control (Fig 7).

## Discussion

Achieving natural regeneration or even establishing planted seedlings in degraded desert lands is extremely challenging, particularly in areas with highly disturbed soils, unprotected environments, and poor physicochemical properties, alongside diminishing conditions of their functional microbial communities. The current research evaluated the influence of inoculating indigenous free-living diazotrophs and root-associated plant growth-promoting bacteria on the growth performance and nutrient uptake ability of four native plant species. The indigenous bioinoculants used in this study were isolated from the rhizosphere soils associated with three native keystone plant species, along with rhizobacteria obtained from *Vachellia pachyceras* root nodules in earlier experiment [20]. Overall our findings, demonstrate that inoculation of indigenous bacterial strains across a range of inoculum densities significantly improved nutrient uptake in all four native plant species grown under controlled greenhouse conditions. These improvements were accompanied by corresponding increase in biomass production, indicating that the inoculation treatment contributed to plant growth. However, the magnitude, direction, and specificity of the responses depend among plant species, inoculum density, and growth medium, highlighting the importance of optimizing inoculum characteristics for different revegetation context Several studies have isolated and characterized beneficial microbes, including mycorrhizal fungi and plant-growth promoting rhizibacteria from arid and semi-arid soils; most of these works have remained descriptive or focused largely on agricultural crop species [22, 28, 29, 30]. However, information on the practical application of such beneficial microorganisms to enhance the growth and nutrient acquisition of native desert plants in the Middle Eastern regions remains limited [5, 20]. The current study contributes to filling this gap by demonstrating the potential of locally adapted microbial inoculants to improve the early establishment of native plant species, offering a promising strategy for restoring degraded desert landscapes.

The significant (p ≤ 0.001) biomass responses observed at both inoculum densities (10^4^ CFU/ml and 10^8^ CFU/ml) indicate that *R. epapposum* responded positively to inoculation, likely through enhanced nitrogen acquisition and possibly improved root system functioning or effective nutrient mobilization. Interestingly, the difference between the two inoculum densities was not significant for either root or shoot biomass, although total plant biomass for the 10^8^ CFU/ml treatment exceeded that of the 10^4^ CFU/ml treatment by 54%. These results indicate that once a functionally effective threshold density is reached, additional increases in inoculum load do not confer further benefits in total biomass growth in *R. epapposum*. In the case of *Farsetia aegyptia*, a similar pattern of enhanced growth response was observed, with markedly greater biomass accumulation, particularly at the higher inoculum density (10^8^ CFU/ml), resulting in a 177% to 182% increase in shoot and total biomass compared to non-inoculated controls (p ≤ 0.001). Similarly, *H. salicornicum* exhibited a substantial increase in seedling biomass at both inoculum densities, with total biomass increasing by 123% compared to non-inoculated controls (p ≤ 0.001). Collectively, these results demonstrate that native desert plant species in Kuwait are highly responsive to inoculation with indigenous bacterial consortia. The observed increases in biomass across multiple species emphasize the potential of regionally adapted microbial inoculants to improve early seedling performance and promote growth under nutrient-limited desert conditions.

*Vachellia pachyceras* was examined across four inoculum densities (10^2^, 10^4,^ 10^6,^ and 10^8^ CFU/ml). Interestingly, in the case of *V. pachyceras* in the Fabaceae family, seedlings inoculated with low to medium densities (10^2^ CFU/ml and 10^4^ CFU/ml) exhibited significantly greater increase in root, shoot, and total biomass, as well as enhanced nodulation (nodule number and dry weight) compared to non-inoculated controls. In contrast, the highest inoculum density (10^8^ CFU/ml) did not produce comparable improvements in growth or nodulation. The highest increases in root, shoot, and total biomass were observed by 190%, 124%, and 154%, respectively, under the lower inoculum concentrations.

The lower response at the highest density (10^8^ CFU/ml) implies potential negative feedback, and this observation suggests a possible saturation effect at higher bacterial concentrations. Excessive cell concentration may limit nodulation efficiency and colonization due to oxygen constrains, resource competition, or interference among microbial populations. Biological nitrogen fixation is an energetically expensive process in which atmospheric N_2_ is converted into ammonia utilizing ATP, which requires carbon investment from the host plant [31]. The high inoculum densities may impose a carbon burden that outweighs the benefits of additional bacterial cells. This may explain reduced growth response in *Valhalla sp*. at 10^8^ CFU/ml treatment level.

These findings are consistent with previous observations reporting density-dependent outcomes in legume-rhizobia symbioses [15, 28, 32, 33, 34], reinforcing the significance of plants optimizing inoculum density to achieve maximum plant benefit. In the context of restoration and revegetation programs, the results indicate that moderate inoculum densities are sufficient to enhance seedling establishment while maintaining cost-effectiveness.

In our experiment, the growth medium substantially affected root biomass across the evaluated species, with significant differences observed in *R. epapposum* (p ≤ 0.011) and *H. salicornicum* (p ≤ 0.007). The magnitude of biomass enhancement in certain species was particularly noteworthy. Increases in root systems are crucial in arid environments, where a well-developed root system enhances the plant’s ability to access limited water and nutrient resources. In our study, the increase in average root biomass by 88% in *R. epapposum* when grown in desert soils compared to potting mix soils, while *H. salicornicum* showed a remarkable 476% increase under the same comparison. These results indicate a strong species-dependent response, and also highlights the influence of substrate quality, environment, nutrient availability, soil structure, and species native ecological growing conditions, which may play a key role in enabling root growth, proliferation, and nutrient uptake, and the performance of microbial inoculation. For *H. salicornicum* and *F. aegyptia*, desert soil supported superior growth, likely reflecting their evolutionary adaptation to native soil conditions and more compatible interactions with indigenous microbial communities.

However, effective plant growth enhancement also depends on successful root colonization by inoculated rhizobacteria, a process that is strongly shaped by the particular ecological niche and the degree of co-adaptation between host and microbial partners [35, 36].

Our results suggest that the success of inoculation may depend on a conducive soil environment that allows roots to colonize and support bacterial symbionts. Pankievicz et al. [31] indicated that effective symbiosis depends on the compatibility between a suitable host plant and diazotrophic bacteria, which is associated with favorable environmental conditions that support optimal nitrogen fixation. Inoculation with indigenous inoculum obtained from rhizospheric desert soil significantly improved plant growth, demonstrating that isolated native bacteria are better adapted to arid soil conditions and remain effective in low-nutrient environments, as observed in a greenhouse experiment.

Across all species, bacterial inoculation significantly increased plant nutrient content, particularly N, P, K, and key micronutrients (Mg, Ca, Na, and S). These increases likely reflect both direct microbial contributions to nutrient supply and indirect effects, including improved root development and i rhizosphere nutrient mobilization. Our study results are in agreement with previous findings showing that plant growth-promoting rhizobacteria enhance growth and nutrient uptake of maize plants when inoculated under greenhouse conditions [37, 38], and improved plant establishment of native plants and soil quality in degraded forest and desert soils [15, 16, 17]. Similarly, in a recent report demonstrated that isolated nitrogen-fixing bacteria from giant reed, and switch grass have the potential to influence plant growth and total nutrient uptake when inoculated to agricultural crops [39]. Similarly, PGPR-based inoculants and biostumulants have been reported to improve wheat growth [40], crop yield and nutrient uptake [41], and N, P, and K uptake of maize grown in nutrient-deficient calcisol soil [28]. Tsegaye et al. [29] further demonstrated that single and consortium PGPR treatments significantly improved teff growth, grain yield and uptake of N, P, K, Ca, and S. Consistence with Egamberdiyeva [28] and Xu et al. [39], our findings confirm that inoculation enhances not only macronutrient uptake but also the accumulation of micronutrients Mg, Na, Ca, C, and trace elements.

In *R. epapposum*, inoculation significantly (P≤0.001) increased N, P, and K uptake by 176-319%, 185-244%, and 215-358%, respectively (*P* ≤ 0.001), with the highest inoculum density yielding the greatest response. Neither growth medium nor its interaction with inoculation influenced nutrient uptake (Table 2), indicating that the bacterial effect was consistent across the soil typess. *F. aegyptia,* exhibited similar inoculation-induced increases in nutrient uptake as observed for increased biomass production. Moreover, in this species, growth medium significantly influenced N, K, Ca, and S uptake, indicating that the soil’s inherent nutrient or physiochemical traits and compatibility with indigenous microbial communities influenced the plant’s nutritional responses. The superior inoculation effect at 10^8^ CFU/ml treatment implies that a sufficiently high bacterial load may be necessary to overcome competition or establishment barriers. These results suggest that optimizing inoculum density and matching soil conditions are critical for maximizing benefits in *F. aegyptia.* These outcomes suggest that the isolates possess full or partial symbiotic capabilities—either forming functional nodules and contributing to N fixation or promoting growth through indirect mechanisms, such as phytohormone production, nutrient solubilization, or root architecture modification. The diminished response at high bacterial density may reflect density-dependent regulation, competition, or suppression effects In *H. salicornicum,* inoculation enhanced nutrient uptake significantly by 70-95% for N, 68-115% for P, and 84-148% for K, along with increased assimilation of Mg, ca, Na, C, and S. These broad improvements indicate that inoculation may facilitate mobilization of multiple macro- and micronutrients in nutrient-poor desert soils. The greater response at 10^8^ CFU/ml versus 10^4^ CFU/ml treatment suggests that higher inoculum density is beneficial in this species.

In contrast, *V. pachyceras,* showed maximum nutrient uptake at lower inoculum density (10^2^ - 10^4^ CFU/ml). While inoculation significantly elevated shoot N, P, K, Mg, Na, Ca, C, and S, the highest density (10^8^ CFU/ml) did not significantly increase N. P, or K uptake. The growth medium and its interactions with inoculation did not have a significant effect on nutrient uptake. The responses in *Vachellia* system indicate a more complex symbiotic system in which isolated root-nodulating bacteria establish effective associations with the host plant primarily at low to moderate densities, likely achieving optimal root colonization and balanced plant-microbe interactions. Such responses suggest that these isolates possess full or partial symbiotic capabilities either forming functional nodules and contributing directly to nitrogen fixation or promoting plant growth via indirect mechanisms such as phytohormore production, nutrient solubilization, or root architecture modifications. The lack of positive response at higher bacterial density may reflect a possible density–dependent regulation, competition, or suppression. Both commercial and indigenous rhizobial inoculants significantly enhanced the growth and nutrient uptake of *V. pachyceras* seedlings under greenhouse conditions. Inoculation with commercial strains of *Rhizobium leguminosarum* ATCC and *Bradyrhizobium* sp. ATCC increased shoot biomass and nitrogen content compared to non-inoculated controls, with *R. leguminosarum* ATCC showing the greatest response (Table 3), indicating the formation of effective symbioses and high N-fixation potential. The growth medium further influenced plant performance, as seedlings grown in potting soil exhibited greater height, shoot biomass, nodule number, and nitrogen content than those grown in desert soil, likely reflecting enhanced nutrient availability and more favorable physicochemical conditions. While greenhouse results demonstrate the promise of these commercial inoculants, field validation is necessary to determine their effectiveness under natural arid-land conditions. Overall, this greenhouse inoculation study confirms that inoculation with nitrogen-fixing bacteria can substantially enhance the growth and nutrient acquisition of desert native plants in nutrient-poor soils, as well as in nursery substrates. These findings are consistent with previous studies showing microbial inoculants increase not only N but also P, K, and micronutrient accumulation, likely through mechanisms such as improved root architecture, increased root exudation, nutrient solubilization, and rhizosphere activation, and phytohormone production [42, 43, 44, 45]. Although the specific mechanisms operating in this study cannot be conclusively identified within the scope of this study, previous studies attribute similar improvements to plant growth promotion, biological N fixation, organic compounds production, disease suppression, and root elongation [28, 46]. Other studies have reported that PGPB can suppress soil-borne pathogens and improve soil structure and microbial diversity, thereby indirectly supporting nutrient absorption and seedling establishment [18, 19]. Increased total nutrient uptake may also reflect increased shoot and root biomass as observed in agricultural crops [47, 48]. To our knowledge, this is the first study to evaluate growth response and nutrient mobilization of *V. pachyceras* seedlings inoculated with potential nitrogen-fixing rhizobacteria from the Kuwait desert, highlighting their potential application as biofertilizers for native plants establishment.

## Conclusion

In general, both indigenous and commercial rhizobial inoculants seedling growth and nutrient uptake compared to non-inoculated seedlings under the current experimental conditions. The study evaluated the collective effects of isolated N-fixing bacteria rather than individual strains, demonstrating that growth improvements, successful nodulation, and N-fixation depend on host-microbe compatibility, inoculum density, and species-specific responses. The inoculation protocol proved effective across the tested native species, highlighting the potential of these isolates to improve growth and nutrient acquisition in Kuwait’s desert flora. While the controlled nursery conditions offer important insights into symbiotic performance, they cannot fully capture the complexity of arid field environments, including heterogeneous soils, resident microbial competition, and environmental stresses. Therefore, field trials are required to validate the performance, ecological adaptability, and nitrogen-fixation efficiency of indigenous and commercial inoculants. Such research will support the development of optimized, field-ready bioinoculation strategies for sustainable restoration and revegetation efforts in desert ecosystems.

## Acknowledgement

The authors immensely acknowledge the Kuwait Institute for Scientific Research (KISR) and Kuwait Foundation for the Advancement of Sciences (KFAS) for the constant support and encouragement throughout the project. We also thank the greenhouse helpers for their assistance in the maintenance of greenhouse experiments and laboratory helpers for laboratory analysis.

## Author Contributions

Conceptualization: M. K. Suleiman, A. M. Quoreshi.

Formal analysis-laboratory and greenhouse: A. J. Manuvel, M. T. Sivadasan.

Statistical analysis: Sheena Jacob

Funding acquisition: M. K. Suleiman, A. M. Quoreshi

Investigation: M. K. Suleiman, A. M. Quoreshi, A. J. Manuvel, M. T. Sivadasan, Sheena Jacob

Methodology: M. K. Suleiman, A. M. Quoreshi, A. J. Manuvel, M. T. Sivadasan.

Project administration: M. K. Suleiman. Supervision: M. K. Suleiman, A. M. Quoreshi. Validation: M. K. Suleiman, A. M. Quoreshi.

Writing – original draft: M. K. Suleiman, A. M. Quoreshi, A. J. Manuvel.

Writing – review & editing: M. K. Suleiman, A. M. Quoreshi, A. J. Manuvel, M. T. Sivadasan, Sheena Jacob.

## Notes

### Competing Interest Statement

The authors have declared no competing interest.

